# Structural variation across 138,134 samples in the TOPMed consortium

**DOI:** 10.1101/2023.01.25.525428

**Authors:** Goo Jun, Adam C English, Ginger A Metcalf, Jianzhi Yang, Mark JP Chaisson, Nathan Pankratz, Vipin K Menon, William J Salerno, Olga Krasheninina, Albert V Smith, John A Lane, Tom Blackwell, Hyun Min Kang, Sejal Salvi, Qingchang Meng, Hua Shen, Divya Pasham, Sravya Bhamidipati, Kavya Kottapalli, Donna K. Arnett, Allison Ashley-Koch, Paul L. Auer, Kathleen M Beutel, Joshua C. Bis, John Blangero, Donald W. Bowden, Jennifer A. Brody, Brian E. Cade, Yii-Der Ida Chen, Michael H. Cho, Joanne E. Curran, Myriam Fornage, Barry I. Freedman, Tasha Fingerlin, Bruce D. Gelb, Lifang Hou, Yi-Jen Hung, John P Kane, Robert Kaplan, Wonji Kim, Ruth J.F. Loos, Gregory M Marcus, Rasika A. Mathias, Stephen T. McGarvey, Courtney Montgomery, Take Naseri, S. Mehdi Nouraie, Michael H. Preuss, Nicholette D. Palmer, Patricia A. Peyser, Laura M. Raffield, Aakrosh Ratan, Susan Redline, Sefuiva Reupena, Jerome I. Rotter, Stephen S. Rich, Michiel Rienstra, Ingo Ruczinski, Vijay G. Sankaran, David A. Schwartz, Christine E. Seidman, Jonathan G. Seidman, Edwin K. Silverman, Jennifer A. Smith, Adrienne Stilp, Kent D. Taylor, Marilyn J. Telen, Scott T. Weiss, L. Keoki Williams, Baojun Wu, Lisa R. Yanek, Yingze Zhang, Jessica Lasky-Su, Marie Claude Gingras, Susan K. Dutcher, Evan E. Eichler, Stacey Gabriel, Soren Germer, Ryan Kim, Karine A. Viaud-Martinez, Deborah A. Nickerson, NHLBI Trans-Omics for Precision Medicine (TOPMed) Consortium, James Luo, Alex Reiner, Richard A Gibbs, Eric Boerwinkle, Goncalo Abecasis, Fritz J Sedlazeck

## Abstract

Ever larger Structural Variant (SV) catalogs highlighting the diversity within and between populations help researchers better understand the links between SVs and disease. The identification of SVs from DNA sequence data is non-trivial and requires a balance between comprehensiveness and precision. Here we present a catalog of 355,667 SVs (59.34% novel) across autosomes and the X chromosome (50bp+) from 138,134 individuals in the diverse TOPMed consortium. We describe our methodologies for SV inference resulting in high variant quality and >90% allele concordance compared to long-read de-novo assemblies of well-characterized control samples. We demonstrate utility through significant associations between SVs and important various cardio-metabolic and hemotologic traits. We have identified 690 SV hotspots and deserts and those that potentially impact the regulation of medically relevant genes. This catalog characterizes SVs across multiple populations and will serve as a valuable tool to understand the impact of SV on disease development and progression.

## Main

A fundamental goal of human genetics is to discover the genetic etiology underlying disorders that plague human health and to build comprehensive models for disease biology. While we know that all classes of variant alleles are impactful^1–4^, structural variants (SVs) have not been characterized as extensively as single nucleotide variants (SNVs), particularly in diverse populations^5^. Structural variants are important types of genomic alterations that are typically described as insertions, deletions, or other complex rearrangements. Structural variants have been shown to play an important role in population diversity^6,7^ and evolution^8–10^. For example, multiple speciation events are reportedly due to de novo deletions and duplications^11^. SVs may also be medically relevant and have been shown to play a key role in cancer^12,13^ and neurological^14,15^ and cardiovascular^16,17^ conditions. Further, their impact in Mendelian disorders is also well documented^18,19^.

Despite their biological importance, our knowledge of SVs is limited^4,20^. Accurately resolving classes of variation that are larger than the length of the individual DNA sequence reads poses challenges^4,5,21^. Additionally, accurate assessment of allele frequencies presents difficulties due to the computational expense and limited availability of highly curated SV data reference resources^22^. Previous projects have contributed to our knowledge of SV frequency across different populations and illustrate the role of SVs in disease, but also demonstrate the limitations of current analytical methods. For example, the 1000 Genomes Project (1KGP)^6,22^ leveraged 2,500 diverse whole-genome shotgun (WGS) datasets at different coverage levels to generate an SV catalog that showed stratification by ancestry. 1KGP highlighted the remarkable variety and complexity of structural variants, but the number of samples and the computational methods were limited. More recently gnomAD SV^23^ provided an important resource across 14,891 samples (10,738 unrelated individuals). Approximately 356,000 SVs were identified across multiple individuals, serving as a valuable resource for allele frequency and disease annotation information. A recent publication from NHGRI’s CCDG program^16,17^ also provided a population-scale map from 17,795 individuals and estimated the relative burden of rare deleterious SVs^17^. Another study from UKBB further highlighted the importance of SV^24^. Despite these advances, the methods for identifying and characterizing complex SVs are still evolving and sample sizes have not been adequate to capture many rare events. Additionally, many ancestral backgrounds remain underrepresented in existing resources.

Joint calling of data from large cohorts can allow for greater sensitivity of detection of rare events and makes it possible to call genotypes on all samples in a cohort, but the computational burden can be prohibitive. Additionally, joint calling must take into account the inferential nature of individual SV detection to maintain sensitivity while also limiting false positives as the size of the sample set increases. These limitations must therefore be considered and a robust QC process must be in place when creating a large population-wide SV call set in order to accurately assess its impact on human health and disease.

We analyzed short read whole genome sequence data from the National Heart, Lung, and Blood Institute’s Trans-Omics for Precision Medicine (TOPMed) program to create the largest, most diverse population-based SV catalog to date using scalable and accurate discovery methods. We describe the methods used to generate this resource and establish the high quality of the data through benchmarking. In addition, we offer key insights on the distribution of variant types observed when analyzing SVs from 138,134 samples. We show the impact of these variants on multiple genes, including many medically relevant regions, highlighting the value of this resource for both studying diversity of the human population and improving our understanding of the role of SVs in disease. Furthermore, we are able to confidently identify low frequency SVs. The TOPMed SV catalog thus provides a high-quality resource representing a wide range of SV frequencies from diverse populations.

## Results

TOPMed seeks to identify factors that increase or decrease the risk of heart, lung, blood, and sleep disorders^25^ by coupling multiple-omic datasets with clinical phenotypes. The primary data includes whole-genome sequencing on 138,134 individuals from 43 studies^25^. Samples were sequenced at an average of 30X coverage using paired-end Illumina technology at five sequencing centers across multiple years^25^. Data was centralized at the TOPMed Informatics Research Center for QC evaluation and initial alignment to the GRCh38 reference. To ensure accurate variant discovery, only samples meeting strict data quality requirements were included in this call set (>30x coverage and genome coverage metrics averaged 95% ≥ 10x, and 90% ≥ 20x coverage). **Figure 1A** shows the composition of the samples with respect to genetically inferred ancestry^25^. While the majority of the samples are of European descent (57.8%), there is substantial representation from African (30.2%) and Asian (7.3%) descent. Other traditionally underrepresented groups make up these general categories. For example, a subset of individuals (n=1,267) are from a Samoan^26^ cohort and the majority appear in the East Asian and Samoan (EAS) category when grouped by genetically inferred ancestry.

**Figure 1:**
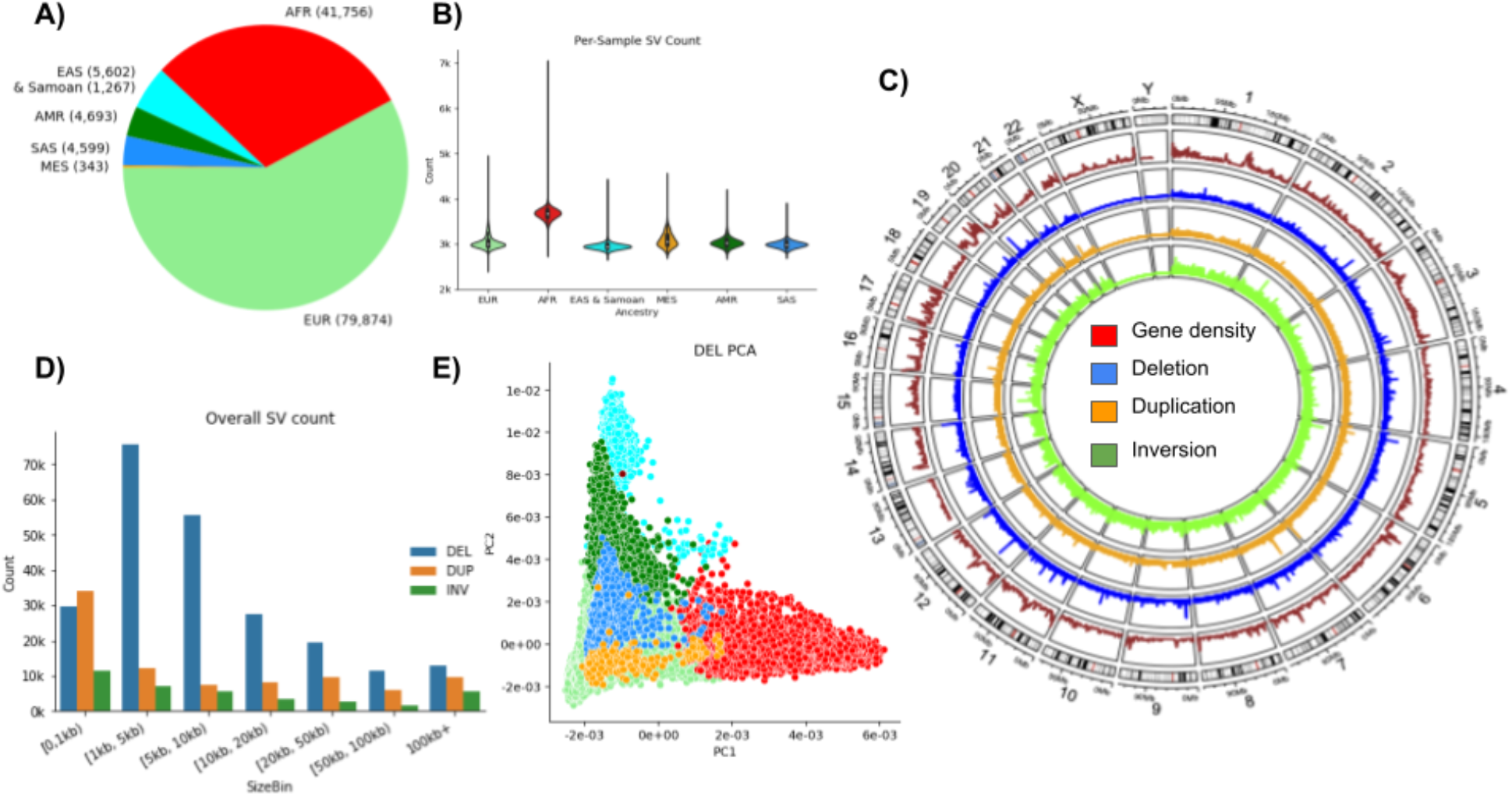
Overview of SV calls. A) Sample counts based on genetically inferred ancestry showing the majority of individuals are Europeans (EUR) followed by African (AFR), East Asian and Samoan (EAS), American (AMR), South Asian (SAS) and Mestizo (MES) ancestry. B) Per-sample SV count distributions by ancestry. C) Overview of gene density (red), deletions (blue), duplications (orange) and inversions (green). D) Size distribution of population genotyped CNV and inversions. The majority of SVs across the population are large events. E) Randomized PCA principal components 1 and 2 of deletions. See supplementary Figures 2-4 for deletions, duplications and inversions.

We implemented a novel SV discovery pipeline to address the computational challenges posed by mega-scale sequencing projects (see Methods for details). In short, each sample was independently analyzed for discordant paired-end, split read, and read depth SV signals using Parliament2^27^. Parliament2 deploys a multi-caller strategy to generate a VCF file of SVs per sample and operates the key principle to leverage the different heuristics that individual state-of-the-art SV callers utilize. Each SV caller has slightly different strengths and weaknesses which are optimized in our SV merging tool, SURVIVOR^10^. The Parliament2 pipeline is optimized for cost-efficient cloud computing and has a high accuracy (F1 score 74.7%) when benchmarked against Genome in a Bottle Consortium (GIAB) standards^27,28^. After the initial calling, we used SURVIVOR merge^10^ to obtain a collection of candidate SVs across all samples. We used muCNV^29^ for efficient population-wide joint genotyping of the candidate SV. The joint genotyping step includes integrating all supporting evidence for each candidate SV across all samples and fitting mixture models that separate carriers of the SV from the rest of the samples. Some samples with low-quality indicators from SNP-based evaluations showed unusually high numbers of duplications and thus were removed as they affected the overall joint genotyping processes. Joint genotyping was performed with stringent quality evaluation by muCNV to produce an accurate set of SV genotypes that can be directly used for genotype-phenotype analyses. We utilized read mapping information and multi-sample statistics to achieve inversion genotyping at scale.

### Summary properties of SVs across 138,134 individuals

**Figure 1B** shows the average number of SVs per individual and across ancestry groups. As expected, individuals of African ancestry have more SVs than those of other ancestries. We also observed a difference for Mestizo (MES)-based ancestries compared to Europeans or Asians. The SV catalog includes a total of 355,667 SVs: 231,817 deletions (65.1%); 86,911 duplications (24.4%); 36,939 inversions (10.5%) in autosomes. **Figure 1C** gives an overview along the entire genome of the SV densities with respect to the gene density. We observe multiple locations with concentrated SV density/hotspots. **Figure 1D** shows the size distribution of the SVs and their types included in this call set. Half of the SVs are at least 5.7 kbp long, with an average SV length of 52.7 kbp (**Supplementary Table 1**). **Figure 1E** shows the principal component analysis plots across deletions. **Supplementary Figure 2-4** shows the PCA analysis for Deletions, Duplications and inversions respectively. We note the expected distribution with stratification being driven by sample ancestry with clustering of samples of African descent (in red) occurring away from the East Asian and European ancestry samples. This can be seen by deletions and duplications independently. However, inversions appear to have an undefined pattern potentially because of the low number of inversions across the SV call set. We note the use of ancestral groupings in this analysis may be limiting due to lack of granularity. For example, a subset of participants from a Samoan cohort are grouped with the East Asian & Samoan (EAS) population group category; and population-specific analysis might reveal more differences for such isolated populations.

The observed allele frequency distribution is consistent with an exponential growth of the human population with 95.5% categorized as rare (<1%) and 47.3% as singletons. The majority of singletons are deletions (101,346) while inversions represent the minority (21,797). Over the entire frequency spectrum (**Supplementary Figure 1A**) we see a small increase of fixed SVs (95-100% AF).

### Establishing a highly concordant and accurate SV call set

We included two control subjects from 1KGP (NA12878 of Central European ancestry, and NA19238 of Yoruban ancestry) in our callset and subsequent analyses. These samples served as process controls for the TOPMed program and were sequenced ~32 times each across each contributing genome center^25^. Furthermore, there are high-quality haplotype-resolved assemblies of these genomes that provide a reference for variant benchmarking^30^. We used two different SV benchmarking methods: TT-Mars^31^ and Truvari^32^ (see Methods for details) to evaluate our variant call sets. On average across the two subjects, there were 3149±32 / 3749±43 deletions, and 200±11 / 279±12 duplications discovered by each sequencing run, indicating a high degree of consistency. The assemblies enabled the assessment of, on average, 93% of calls (**Figure 2A)**. The positive predictive value (PPV) estimated by TT-Mars averaged 0.90 for NA12878 deletion calls and 0.87 for NA19238 deletion calls (**Figure 2B**). The estimated PPV based on Truvari analysis indicated that 87% of deletions were true positives. Variant calls for NA19238 had similar (though improved) metrics of 95% (TT-Mars) and 89% (Truvari) PPV. The combined assessments of TT-Mars and Truvari allow us to estimate the overall precision of our deletions to be 90.0-91.5%. We could assess between 96 and 98% of all duplications. The overall PPV of duplications was determined to be 86.5% by TT-Mars and Truvari (**Figure 2A & B**)^28^. In addition, we reviewed SVs for NA12878 that were not confirmed by either method (n=219) using Samplot^33^. In most cases, there was clear read evidence that there was a real CNV at the predicted location (61% of variants that were initially non-confirmed by TT-Mars or Truvari). The others that were non-confirmed appeared to be a result of inaccurate breakpoints. This was pronounced in regions of low map-ability/ repetitive regions and explains why both methods(TT-Mars or Truvari) did not confirm these calls. We were not able to establish visual evidence for 6% of non-confirmed SVs to confirm their presence in these control samples.

**Figure 2:**
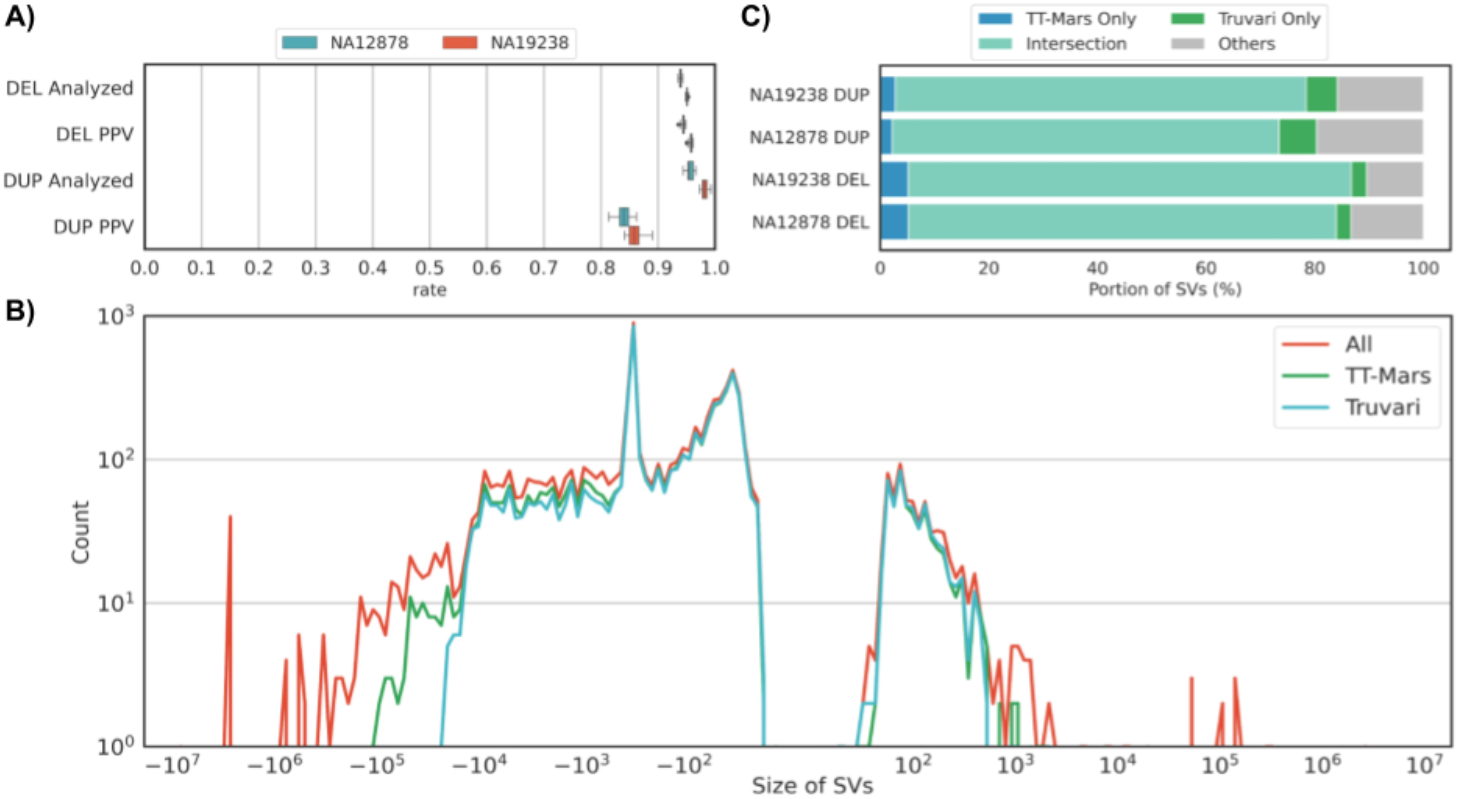
Evaluation of SV call sets against using haplotype-resolved assemblies for deletion (DEL) and duplication (DUP) calls in the NA12878 and NA19238 genomes. A) Evaluation using TT-Mars. The fraction of calls that may be assessed using the assemblies (analyzed) and positive predictive value (PPV) are given for DEL and DUP calls. B) Support for calls from both the TT-Mars and Truvari methods. C) The size by count spectrum of all calls (red), the count validated by TT-Mars (green), and the count validated by Truvari (blue) for the combination of both genomes.

To estimate consistency, we counted the SVs per subject that were assessed as being a true positive in at least one replicate and measured the percentage of these SVs that each replicate recovered. For NA12878, Truvari found 3,004 DELs and 282 DUPs, 93% and 59% of which were recovered per-replicate on average, respectively. For NA19238, the 3,906 DELs and 411 DUPs were recovered in 87% and 63% of replicates, respectively. TT-Mars and Truvari use complementary approaches for variant validation and so it is interesting, but perhaps not surprising, that each method highlights certain deletions or duplications that can be verified by only one of the methods (**Figure 2B**). Neither validation approach confirmed deletions greater than 500kb or duplications over 50kb (**Figure 2C)**. Deletions and duplications of less than 1kb account for 78% and 95% of calls, respectively, and the PPV in this size range was 93% and 87%, indicating a high accuracy for shorter calls.

We evaluated the genotyping accuracy by measuring error rates from Mendelian consistencies using 11,387 samples in 4,081 trios and 173 duos (**Figure 3A**). The estimated error rates are 0.29% for deletions, 6.1% for duplications, and 6.1% for inversions. Inheritance patterns indicate de novo variants at 0.33% of heterozygous deletions, 1.4% duplications, and 9.0% inversions. Mendelian inconsistencies and de novo variant rates are comparable to each other in deletions and inversions because they are all biallelic variations, but the Mendelian error rate is much higher than the de novo error rate in duplications due to multi-allelic variations. The allelic balances (**Figure 3A)** were 49.6:50.4 (REF:HET) in deletions, 55.2:44.8 in duplications, and 57.9:42.0 in inversions. Compared to the expected 50:50 distribution, we see slightly more REF calls than HET calls in duplications, meaning there exists a slight reference bias. Overall, the ratios do not show excessive reference biases and do not deviate significantly from the expected. The average het/hom ratio of samples is 2.26±0.34 (**Supplementary Table 2**)

**Figure 3:**
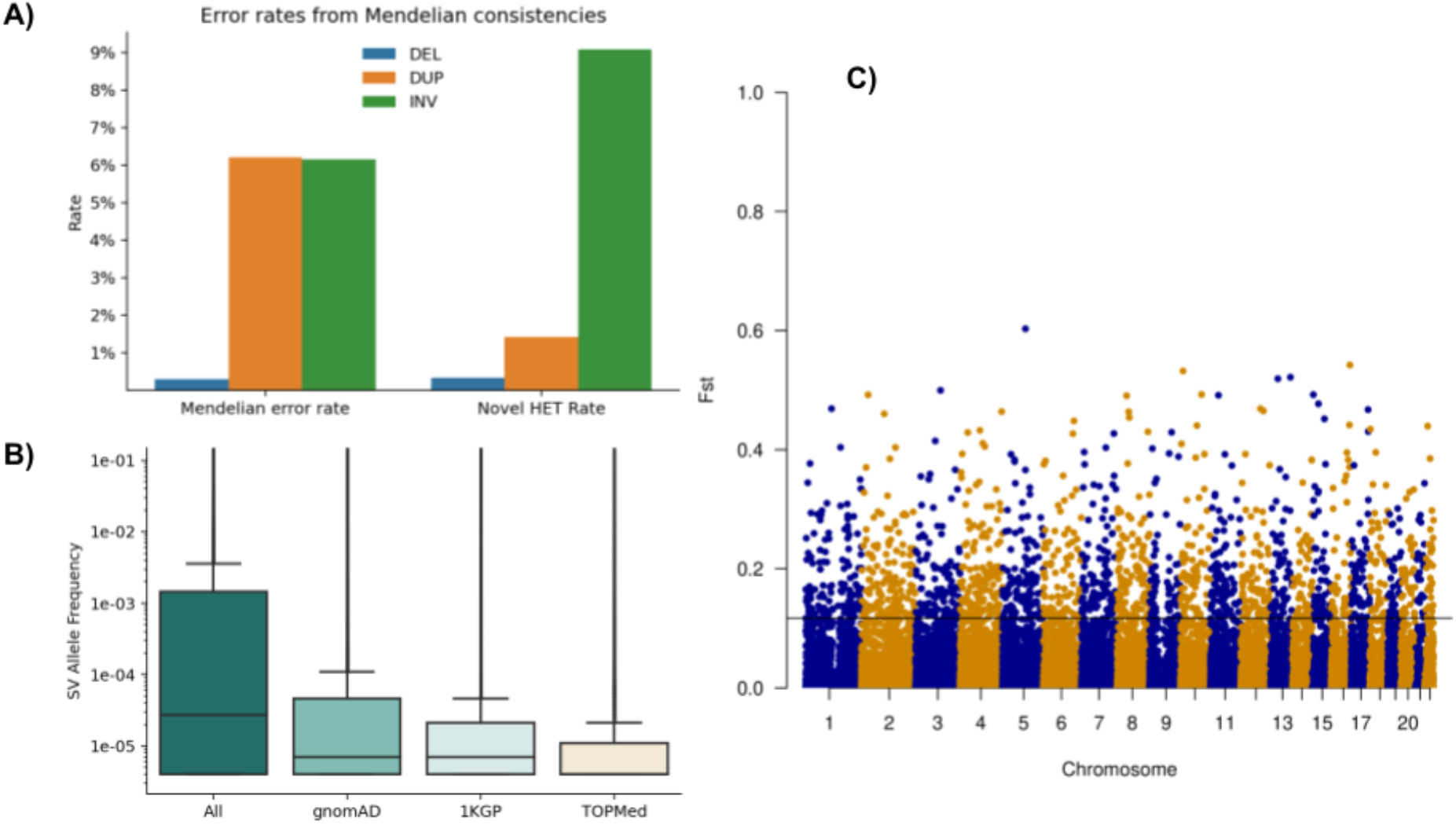
A) Assessment of Mendelian error and novel HET rates per SV type across 11,387 samples from trio/duo families. For the evaluation, we also include the multiallelic variants that do increase the error rate, especially across duplications. B) Overlap of SV over 1KGP and gnomAD SV with respect to the allele frequency within the TOPMed SV call set. The allele frequencies change slightly between the novel SV from TOPMed and other overlapping SV. C) F_ST_ plot of African versus European ancestry of SV across the entire genome, highlighting a threshold of 0.11.

### Identification of 163,794 novel SVs

To evaluate the novelty of the TOPMed SV call set, we compared it with the gnomAD SV (14,891 samples) and 1KGP (2,506 samples) call sets, which capture variants with allele frequency down to 0.01%, and 0.1%, respectively. Events identified across callsets are classified as the same SV if they have an overlap of ≥70% using AnnotSV^34^. Across all types of SVs, 52% of TOPMed SVs have overlap with gnomAD SV or 1KGP, with the highest overlap for deletions (56.8%) followed by duplications (48.0%) and inversions (46.1%). As expected, we observed a larger overlap with gnomAD SV (44.05% of calls) compared to 1KGP (28.50%). The variants unique to TOPMed are consistent with the number of rare events detected using increased sample sizes. **Figure 3B** shows the overlap of TOPMed SV to 1KGP and gnomAD SV with respect to the allele frequency of the variants. Events found in all three catalogs have the highest mean allele frequency (1.9%) and events exclusively in TOPMed have the lowest (0.4%). Our call set also includes 168,307 singletons (47.28% of calls), 45% of which overlap with a gnomAD or 1KGP call. For variants shared across individuals (AF≥0.01) and overlapping between TOPMed SV and gnomAD, the Pearson correlation coefficient between reported allele frequencies is 0.85.

### SV association with human phenotypes and further applications

The availability of a validated SV catalog on a large diverse population-based sample set creates an important resource for genome-wide association analyses between SVs and a broad array of clinically relevant phenotypes. As a proof-of-concept for the SV-phenotype association analyses, we tested genome-wide association between SVs and various cardio-metabolic phenotypes from Atherosclerosis Risk in Communities (ARIC) and Hispanic Community Health Study / Study of Latinos (HCHS/SOL) cohorts. In ARIC, we had 3,642 samples from European American ancestry and 269 samples from African American ancestry with SV genotypes. In HCHS/SOL, we had 3,945 samples with SV genotypes. We tested for association in each ancestry group separately using EPACTS (ARIC AA/EA) and GMMAT (SOL) and performed a meta-analysis across three ancestry groups using METAL. Although we tested only a modest sample size compared to the entire TOPMed set, we were able to successfully identify genome-wide significant associations. We identified a 3.8 kbp deletion overlapping *HBA2* in human alpha globin cluster (chr16, 173,578-177,341) that was significantly associated with decreased hemoglobin (*p*=7.93×10^−15^, standardized *β=* −0.34) and a 893 bp deletion overlapping *BCRP3* (chr22, 24,637,046-24,637-939) that was significantly associated with increased gamma-glutamyltransferase (GGT) levels (p=1.94×10^−8^, standardized *β*= 0.18). The *HBA2* deletion was a previously reported deletion and is known to be associated with red blood cell traits and a recent TOPMed-wide association analysis confirmed the association with seven red blood cell related traits^35^. We also genotyped this specific variant across additional ARIC and SOL TOPMed/CCDG WGS samples and identified a significant association with decreased hemoglobin, decreased hematocrit, and increased HbA1c levels^16,36,37^. The association between *BCRP3* deletion and GGT levels has not previously been reported, but is located approximately 9 kb downstream of the *GGT1* gene that directly encodes the GGT enzyme. Interestingly, both deletions were common in AA but very rare in EA (*HBA2* AA MAF=0.20, EA MAF=0.004; *BCRP3* AA MAF=0.51, EA MAF= 0.00077), suggesting the importance of the current multi-population WGS analyses and also the role of SVs that contribute to inter-individual and inter-population differences in the distributions of several clinically relevant biomarkers. Our results show that the call set provides unique value as the largest-scale SV genotype resource available for WGS data and can be utilized for the study of functional contributions of SVs to various complex human traits. Since the release of the SV callset, there has been active involvement of various phenotypic working groups of TOPMed who have reported findings from significant SV-phenotype association; including the hematologic traits^35^ and atrial fibrillation (AF) (manuscript in preparation). The SV set also has been utilized to generate a database of SV-SNP linkage disequilibrium (LD) statistics from subpopulations of TOPMed samples (http://topld.genetics.unc.edu/topld/) ^38^.

### LD between SVs, SNPs, and known GWAS hits

To understand how SVs are associated with other nearby genetic variation, we performed pairwise linkage disequilibrium (LD) analysis between each SV and small variants within a +/- 1 Mbp region around the SV. We used the TOPMed SNV call set freeze 9^25^, and only founder samples were selected from both the small variant and SV call sets. We analyzed 36,056 SVs with MAF>0.001, including 30,191 deletions, 3,717 duplications, and 1,861 inversions. We ran PLINK pairwise LD analyses with a minimum r^2^ value of 0.05. 12,204 SVs had at least one small variant with r^2^>0.3. We identified 28,424 unique small variants with r^2^>0.3; they were distributed most densely in the HLA region of human chromosome 6, a known recombination cold spot.

Another important way to use SNV data in conjunction with SVs is to look at correlation of SVs events with known GWAS signals. Most of the known GWAS association signals are from small variants, but this is also frequently the only variant class evaluated in large association studies. While these GWAS signals may be significant, the functional role of small variants is often obscured. Based on Mendelian disease gene discovery and follow-up studies ^39^, existence of an SV highly correlated with a known GWAS SNP suggests a functional contribution to the associated phenotype. To understand the overall distribution of SVs in the context of reported GWAS signals, we compared data from the NHGRI-EBI GWAS Catalog^40^ to identify SVs in high LD with their reported association signals. From the GWAS Catalog, we identified 201 SNPs with r^2^>0.5 and 132 SNPs with r^2^>0.8 that are associated with at least one SV. For example, a 684-bp deletion in an intergenic region of chromosome 9 (chr9:101,457,417-101,458,101) is in almost perfect LD (r^2^=0.997) with rs2183745 at chr9:101,456,893, which is strongly associated (P=2×10^−133^) with alkaline phosphatase (ALP) level. Another example is a 928-bp deletion upstream of *INSR* at chromosome 19 (chr19:7,258,353-7,259,281) that is in strong LD (r^2^=0.979) with rs12798472 at chr19:7,257,979, which is significantly associated (P=1×10^−58^) with systolic blood pressure. These two SV are in regulatory and intron regions, respectively. The majority of GWAS SNP-linked SV were deletions, but we also found a 55-bp duplication at 19q13.11 that is in strong LD with rs66528626, which is associated with serum albumin levels (P=4.00×10^−9^), and a 200-bp inversion at 10q22.2 that is in strong LD (r^2^=0.953) with rs648078, which (together with a couple of other inversions) is associated with atrial fibrillation (P=6×10^−27^). The full list of SVs in high LD (r^2^>0.8) with SNPs from the GWAS Catalog is in **Supplementary Table 3**.

### Population differentiation of SV

We assessed the extent this dataset contains SVs that exhibit population differentiation measured by Wright’s fixation index F_ST_. There are 2,072 deletions and 21 duplications with F_ST_>0.2 between European (EUR, N=79,874) and African (AA, N=41,756) individuals (**Figure 3C**), the two ancestries that make up the majority of this cohort (**Figure 1A**). Of these high Fst SV, 929 deletions (44.84%) and 14 duplications (66.67%) overlap genes. Because this sample size enables the study of rare SVs, it potentially contains rare events that show population differentiation (excluding singletons). However, among this pair of populations there were no deletion SVs with F_ST_>0.2 with an allele frequency less than 0.05. This is consistent with previous observations that F_ST_ is lower at lower allele frequency^41^ or is dependent on methods of estimation^42^. In addition, we identified 187 genes (**Supplementary Table 4**) with a high F_ST_ from 0.2 to 0.54. Although there is a high correlation between F_ST_ calculated for deletion loci in this sample and F_ST_ calculated for matching variants discovered in the 1000 Genomes Project^6^ (r^2^=0.87), there are 1,063 deletion loci not discovered in this study with F_ST_>0.2 not found in the smaller studies.

### SV annotation and potential impact

To investigate the potential impact of the SVs (see Methods) we used AnnotSV^43^ for the gene annotation, which reports overlapping genes as well as genes directly impacted by the breakpoint of an SV itself (see Methods). Many SVs (47.05%) overlap genes given the nature of our call set (average SV size 52.5 kbp), but we restricted our analysis to common events with an allele frequency above 5%. We identified 2,295 genes overlapping deletions of 5% allele frequency or higher, and 22.18% of these were impacted by more than one structural variant. These genes were significantly enriched across cell junction (FDR: 1.4E-7), ATP binding (FDR: 1.9E-8), and other GO terms and impacted 27 KEGG Pathways (FDR smaller 8.8E-2) with the glutamatergic synapse pathway being the most significant (FDR: 6.3E-4). We investigated the genes impacted by duplications that have an allele frequency of 5% or higher. Across this list, genes with alternative splicing were enriched (FDR: 8.1E-14) as well as zinc finger (FDR:4.1E-7). The genes that were impacted by duplications also showed a significance of the keyword polymorphism (FDR: 6.1E-3) and sequencing variant (1.5E-2). These could again highlight the role of duplications in the promotion of other types of genomic variants. We investigated if these genes overlap any medically relevant genes to determine potential impact on disease. For this, we utilized a list of 5,131 genes previously annotated^44^. Interestingly, 1,351 medically relevant genes were impacted by deletions (AF>5%), which represents a slight majority (58.87%) of all genes impacted by deletions in our call set. For duplications (AF>5%) we identified 801 genes that are impacted and medically relevant. Again, we observed a similar small majority (56.41%) across all genes impacted by these duplications.

We identified SVs that are present in a large portion of the population (AF>90%) and thus likely represent a minor allele in the available reference. For deletions, 113 genes were identified to be potentially impacted. Of these, 65 (57.52%) medically relevant genes are overlapping deletions with AF>90%, which highlights the importance of this SV catalog for ongoing association studies. The TPTE gene was the only one reported to be potentially impacted multiple times, which is a paralog of TPTE2. We identified 55 genes overlapping duplications that are highly shared, meaning that it is likely a copy is deleted on the reference genome. Of these 55 genes, 34 (61.82%) are recorded to be medically relevant in different databases. Two genes GUSBP1 and MUC3A are reported with two highly shared duplications.

Finally, we investigated SVs that occurred in 20 (AF=0.0001) to 100 (AF=0.0007) individuals only. These represent very low allele frequency cases that other studies were potentially under powered to detect. This range includes 13,320 deletions and 10,268 duplications of which 74.40% and 52.28% are present in gnomAD SV, respectively. Across these deletions, we identified 3,360 genes that are potentially impacted. Next, we investigated if these genes are reported to be medically relevant based on an annotated list^44^ and found that 1,884 (56.07%) of these genes are reported to be medically important. The list of enriched GO terms was very diverse as expected, with intracellular signal transduction being the most significant (FDR: 9.7E-4). Interestingly, the number of genes impacted by duplications in this low-frequency spectrum has slightly increased compared to deletions to 3,842 genes. Further, we identified 2,200 (57.26%) genes that are overlapping with previously reported medically relevant genes. Similar to deletions, we also observed intracellular signal transduction being the most significant (FDR: 1.8E-2) GO entry.

A component of the AnnotSV^43^ method is to leverage ClinGen ranking^34,45,46^ annotations of regions with potential haploinsufficiency (HI) and triplosensitivity (TriS) with respect to SVs. HI is the genetic relationship of a loss of one copy introducing a phenotypic effect. TriS is where an additional copy of a gene may lead to a phenotypic effect. **Figure 4A** shows the SV counts by annotation across three allele count bins. In total, 25,645 SVs have HI annotation, within which 3,514 SVs are VUS with 55.29% being deletions and 4,132 SVs pathogenic (50.39% deletions). For TriS, we observed fewer SVs (22,065) with proportions of VUS (366) and pathogenic (17) SVs, likely due to the curation criteria. Of the 4,149 SVs annotated as HI or TriS pathogenic, 85% have an allele count less than 20. SVs associated with Hi and TriS remain to be investigated when combined with traits collected from the TOPMed studies.

### Identification of genomic SV hotspots and deserts across individuals

The large number of individuals in this study gives us the opportunity to further investigate the mutational landscape of SVs across the human population. **Figure 1C** already indicated multiple hotspots or deserts along the genome. To do this more systematically we identified potential hotspots and deserts (0 SV) across the TOPMed SV call set, which represent either higher levels of SVs or potentially conserved regions. We identified hotspots and deserts using a window approach (100 kbp): if the number of SVs is more or less than three times the standard deviation of the mean (see Methods), that window is assigned to be a hotspot or desert, respectively. Using this method we identified 502 deserts and 188 SV hotspots, as shown in **Figure 4 B**. The deserts could represent SV regions that were flagged during QC or ignored by the calling due to repetitiveness or incoherent alignments; to address this question, we compared the locations with the long-read based SV calls from the Human Genome Structural Variation Consortium (HGSVC) from their high-quality assemblies (see Methods). An average of 10.6 TOPMed SV and 1.4 HGSV SV were found in windows with a standard deviation of 12.4 and 2.8, respectively. Doing so identified 226 (45.02%) SV deserts that HGSV and our call set agreed on. Of these 226 candidates, 104 (46.02% of all deserts) overlap 91 genes (**Supplementary Table 5**, see Methods). A standard GO-term analysis did not reveal any significant enrichments. The 91 genes contained 12 long noncoding RNAs, 8 microRNAs, and 7 long intergenic non-protein coding genes. In addition, we also predicted 14 (2.79%) SV desert regions of the genome that disagree with HGSV. This represents only a small fraction of our genomic candidates. Furthermore, we identified 262 (52.19%) deserts that were not classified as deserts or valleys over the analysis of HGSV data. These could be either identified based on the fact that our call set is much larger compared to the 32 genomes that we used here from HGSV, or it could in fact represent additional missing SVs. Overall, our SV desert regions did not show a clear correlation with SNV indicating that an SV desert could still include a multitude of SNV. For hotspots, we only observed an agreement of 4 regions with respect to HGSV data sets. The majority of our predicted hotspots (119, 63.30%) are not classified as either based on the HGSV data set. In contrast, 65 (34.57%) of our predicted hotspots were classified as valleys in HGSV assemblies, which probably highlights lower frequency variations in our call sets. Our SV catalog enables a deeper insight into the regions of the genome that appear ultraconserved (SV deserts) or highly variable for SVs. The latter especially is highly important to consider for SNV and other genetic analyses as they could manifest in incorrectly mapped reads^47^. Further analysis should be conducted to determine if these regions have an impact on phenotype, or if they represent normal genomic variability in a population.

**Figure 4:**
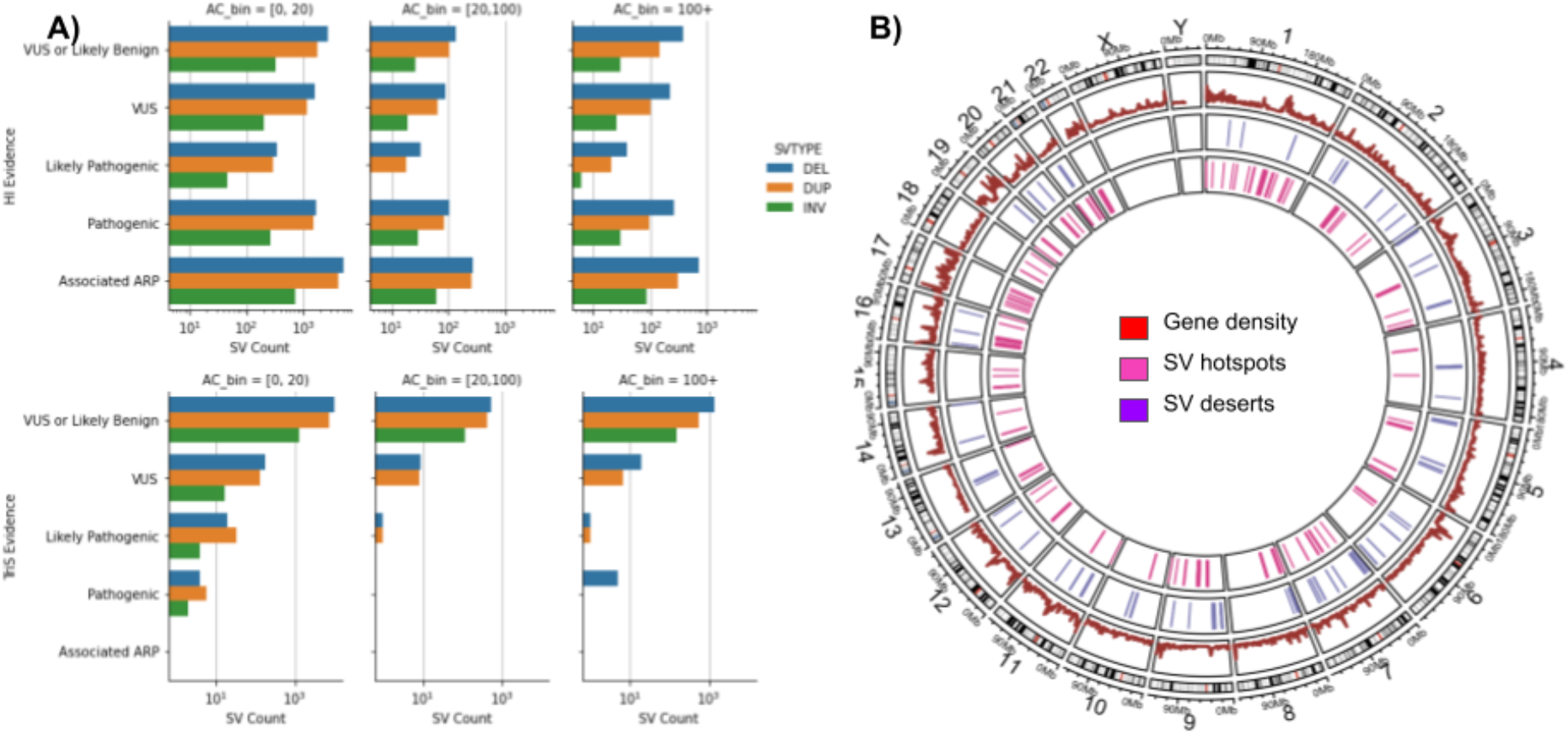
Overview of the impact of the SVs and their clustering along the genome. A) SVs identified and clustered based on their haploinsufficiency (HI) and triplosensitivity (TriS) potential across different allele counts. B) Overview of SV hotspots and deserts across the TOPMed cohort. Here deserts are regions of the genome with no SV identifiable despite the large collection of individuals in this study.

### Structural Variants in chromosome X

The previous analyses and variant counts reflect structural variants in the autosomes only. The nature of the sex chromosomes adds an additional complexity in the interpretation of heterozygous vs. homozygous calls across males and females. The current methods do not allow for accurate genotyping on chromosome Y due to the variation in mapping and read depth and a haplotype based genotyping method is under investigation for application to the Y chromosome. Similar to the autosomal SV calls, we leveraged the two control samples to identify a core set of SVs for studying sex chromosome X specific events. Using this call set we measured an average 0.94 and 0.788 positive predictive value for deletions and duplications, respectively. The two replicate subjects are one male and one female and we observed similar PPV for deletions (0.945 and 0.948) and higher PPV for the female replicate subject’s duplications (0.729; 0.843). We compared the SV calls on chromosome X with those available in gnomAD and 1000 genomes and identified 51.6% of chrX calls that were not previously reported. Separated by project, 22.2% of the TOPMed SVs overlap with 1KGP SVs and 40.5% overlap gnomAD while 14.5% of our chromosome X SVs overlapped with both gnomAD and 1KGP. We identified a significant (p=0.0294, Fisher exact test) difference between known males vs. female SV. **Supplementary Table 6** shows the results of known and novel SV with respect to gender. Overall, there are 8,956 (60.16% novel) SV specific to females and 2,973 (57.89%) SV specific to males.

In total we identified 9,814 deletions, 6,756 duplications and 2,597 inversions across the chromosome X. Per-sample, we find an average of 181 deletions, 12 duplications, and 3 inversions. As expected, we observed 1.5 times more SVs per-sample across the 75,547 (53.54%) female samples of TOPMed compared to the 65,582 male samples, which averages to 235 SVs for females and 148 SVs for males. Furthermore, in females we observed a het/hom ratio of 3.06 for deletions, 3.35 for duplications, and 1.98 for inversions. For homozygous alternative SVs we observe a few more SVs in males (149 SVs) vs. females (58).

## Discussion

The resource described includes 355,667 well-curated structural variants (SVs) from 138,134 diverse human whole genomes (**Figure 1A**). These highly accurate calls coupled with the extensive phenotype data available for the TOPMed cohorts provide an unmatched opportunity to identify associations and ultimately study mechanisms by which these variants impact disease onset and progression. This is already exemplified by Wheeler et all.^35^ who analyzed the SV calls from a subset (N=50,675) of samples for which multiple blood traits had been measured to identify phenotypic associations. They identified 33 independent SVs (23 common and 10 rare) implicated across multiple phenotypes. Interestingly, most of the SVs seem to impact the regulation of genes rather than showing a direct impact on the coding regions^35^. Most SVs were observed around regions of the genome that had previously been identified to play a role in blood traits. However, one SV was shown to impact *KCNJ18*, which represents a novel finding based on this SV catalog. In another instance, Wheeler et al show a ~13kbp deletion (chr7) which includes the EPO promoter and was associated with HGB/HCT trait and in strong LD with a previously reported SNV (rs4729607)^35,48^. The same deletion was also shown in a previous study to impact the expression of genes^2^ including TFR2 and EPHB4 that are involved in iron metabolism and erythropoiesis^49,50^ Thus highlighting the importance of the SV as potentially causative instead of the SNV alone. We anticipate additional disease trait associations as we have identified a large number of genes, including those designated as medically relevant, which contain SVs that are expected to alter expression.

The large sample size across multiple populations and inclusion of low frequency SVs (20-100 individuals, AF 1E-3 to 7E-3) enables exploration of alleles that may be absent from other studies. For example, in 1KGP and gnomAD SV, these low frequency SVs would most often appear as singletons and on average will only be present in 0.25 or 1.4 individuals, respectively. We note that these rare events show an impact across 1,884 medically important genes in the TOPMed call set. This is likely a result of the broad range of disease cohorts that are represented in TOPMed and highlights the diversity across some of these genes. In addition, the large sample size and uniform data quality enabled us to identify different hotspots and deserts of SVs. Despite the large number of individuals, we still identified multiple deserts (regions with no SVs present) and compared them with other studies. Characterization of these patterns can be medically relevant and is important for disease biology, but also may provide valuable insights on human evolution and selection.

The large volume of the TOPMed whole genome sequencing data provides ample opportunity for discovery, however it presents computational and management challenges when generating cohort wide variant resources. To accommodate this scale we implemented the Parliament2 and muCNV framework. Cloud computation was leveraged and the crams were analyzed directly in their native GCP environment, obviating any egress charges. In order to effectively scale to the sample size present in TOPMed, we optimized the runtime of the 5 selected SV callers to fully leverage a 16 core cloud instance. For example, we reduced the wall time of Lumpy from 6.45 to 0.45 hours and for Delly from 8.52 to 0.67 hours. This enabled efficient and accurate processing of individual samples. Despite a high precision rate from Parliament2^27^, processing such a large number of samples results in a high rate of false positives due to the inferential nature of SV callers. To mitigate this we implemented muCNV, which utilizes coverage, split and discordant paired-end reads together to jointly genotype and filter SVs. muCNV achieves a higher precision by computationally validating each SV site across all individuals simultaneously. In addition to muCNV, we leveraged multiple populations and long read-based comparison filtering steps outlined in the methods to achieve a highly accurate SV resource.

It is well established that long read technologies (e.g Pacific Biosciences and Oxford Nanopore) more accurately resolve SVs and have recently shown marked improvement in overall data quality. Despite improvements in quality, price point and capacity, long-read sequencing is still prohibitive for large scale human WGS studies. While there is a natural tension between cost and comprehensiveness, there are opportunities for complementarity where the resolution offered by long read data can be leveraged to improve large scale short read resources. Long read efforts such as GIAB and HGSVC promote SV calling optimizations and the development of novel methodologies to improve the detection of SVs across complex regions in short read data. We demonstrated how long-read control samples from such programs can be used to assess the accuracy of a population-based SV catalog. Nevertheless, by doing so we also discovered slight disagreement between two state-of-the-art benchmark methodologies (Truvari and TTmars). While this might be a minor point of this manuscript, it shows that the SV field has not yet found a standard or agreement of how to compare SV alleles and speaks to the need for continued work in this area.

As a result of this comprehensive analysis of short read sequence data and precise SV calling, this TOPMed resource will provide unparalleled discovery opportunities by presenting allele frequencies across a wide range of individuals and populations. With the ever-increasing number of available Illumina WGS datasets, this call set will provide continued utility in identifying novel associations between complex alleles and disease phenotypes.

## Methods

### SV call set generation

#### Parliament process

For SV calling we deployed our Parliament2^27^ method. Parliament2 processes aligned Illumina paired-end reads and identifies SVs via five programs: Manta^51^, Delly^52^, Lumpy^53^, Breakseq^54^, and CNVnator^55^. Subsequently, we merged the SV calls of the same type using SURVIVOR^10^ merge with a 1 kbp wobble distance between breakpoints. The single sample calling was done in parallel across Google Cloud instances and the results were written to a bucket on DNAnexus. Then another population merge was performed with SURVIVOR merge (1 kbp wobble) and type matching following recommended parameters on a per-chromosome basis. We ignored any translocation calls among the samples. The population VCF file of SV candidates generated by SURVIVOR was passed along for population genotyping.

#### muCNV

The muCNV pipeline is designed for whole-genome population-level joint genotyping of SVs. We generated summary pileups, which recorded all discordant read pairs, split reads, soft clips, average depth information for each candidate SV event, and average depth for each 100-bp interval across the whole genome in a single scan of a CRAM/BAM file^29^. The pileup process also generated per-sample GC curves for GC correction. The pileups (100 to 200MB per sample) were then merged across samples and sliced by chromosomal regions for efficient handling of large sample sizes by a single compute node, as joint genotyping needs concurrent access to pileup data from all samples. Joint genotyping was performed across all samples by combining all supporting information around each candidate SV. To genotype deletions and duplications, we fitted a two-dimensional Gaussian mixture model with 1) the number of supporting alignments (discordant read pairs, split reads, and soft clips) and 2) the normalized read depth. Some deletions and duplications had complex breakpoints, which resulted in a lack of alignment support but clear depth-based signals. These events were genotyped by fitting a mixture model with only read depth information. Inversions were genotyped by fitting a mixture model with only alignment support. We also genotyped candidate SVs with clinical implications as reported in dbVar (https://www.ncbi.nlm.nih.gov/dbvar/studies/nstd102, August 2020 release), which added 453 deletions and 65 duplications.

#### Additional filtering/flagging

We took additional steps to identify potentially low-confidence duplications as they are more sensitive to genomic context and sequencing depth variations. First, we flagged DUPs with aberrant normalized depth as measured by muCNV and highly concordant genotypes with significantly overlapping DUPs. This filter identified and flagged 58,322 DUPs as *PreFiltered*. A second filter involved training a support vector machine (SVM) with five SV features reported by muCNV. These features are mean and standard deviation of the sequencing depths both before (INFO field=PRE) and after (INFO field=POST) the DUP, plus the GC content of the DUP region. Training data were DUPs genotyped as being present in the validation replicate samples and were labeled using Truvari^32^ (see SV validation below). Hyper-parameter tuning of the SVM’s RBF kernel was performed using grid search. An SVM score cutoff was manually set by evaluating increases in PPV versus number of DUPs flagged. As a result, 29,897 DUPs were flagged as being *LowQual*. The SVM was coded using Python version 3.7.12, scikit-learn version 1.0.1, and NumPy version 1.19.5.

#### Chromosome X calling and filtering

Chromosome X has been called separately from autosomes as it involves additional processing for joint calling of males and females. We used inferred sex information based on the ratio between average sequencing depths of chromosomes X and Y and normalized sequencing depth of male samples has been increased by 0.5 before genotyping step. Pseudoautosomal regions (PARs) have been excluded from genotyping. To minimize possible artifacts from depth compensation and mapping issues, we applied SVM-based filtering on all deletions, duplications, and inversions. Features used are mean and standard deviation of sequencing depths, difference in allele frequencies between female and male samples, difference in call rate between male and female samples, and also based on the existence of split-read and soft-clip based support for breakpoints. SVM was trained using variants labeled as true positive or false positive by Truvari^32^. The SVM was coded using Python version 3.7.12, scikit-learn version 1.0.1, and NumPy version 1.19.5.

### SV validation

We used a set of haplotype-resolved long-read assembly^30^. http://ftp.1000genomes.ebi.ac.uk/vol1/ftp/data_collections/HGSVC2/release/v1.0/assemblies/20200628_HHU_assembly-results_CCS_v12/assemblies/phased/v12_NA19238_hgsvc_pbsq2-ccs_1000-pereg.h1-un.racon-p2.fasta http://ftp.1000genomes.ebi.ac.uk/vol1/ftp/data_collections/HGSVC2/release/v1.0/assemblies/20200628_HHU_assembly-results_CCS_v12/assemblies/phased/v12_NA19238_hgsvc_pbsq2-ccs_1000-pereg.h2-un.racon-p2.fasta http://ftp.1000genomes.ebi.ac.uk/vol1/ftp/data_collections/HGSVC2/release/v1.0/assemblies/20200628_HHU_assembly-results_CCS_v12/assemblies/phased/v12_NA12878_giab_pbsq2-ccs_1000-pereg.h1-un.racon-p2.fasta http://ftp.1000genomes.ebi.ac.uk/vol1/ftp/data_collections/HGSVC2/release/v1.0/assemblies/20200628_HHU_assembly-results_CCS_v12/assemblies/phased/v12_NA12878_giab_pbsq2-ccs_1000-pereg.h2-un.racon-p2.fasta

We created a baseline set of SVs per haplotype via Minimap2 v2.17 using Ebert long-read haplotype-resolved assemblies for two subjects, and we used Truvari^32^ collapse v2.1 to perform intra-sample haplotype merging. For each sequenced replicate of the two subjects, we used Truvari v3.1-dev bench to compare variants that were present (i.e., non-reference homozygous and non-missing genotypes) to the baseline variants. Benchmarking comparison parameters were set at {refdist: 500, pctsim: 0.0, buffer: 0.1, pctsize: 0.5, pctovl: 0.0, typeignore: false, use_lev: false, chunksize: 1500, gtcomp: false, sizemin: 10, sizefilt: 10, sizemax: 100000000, passonly: true, no_ref: c, includebed: null, multimatch: true}.

In addition we used TT-Mars ^31^ with the following code:

reference=hg38.no_alts.fasta;output_dir=./; vcf_file=callset.vcf; centro_file=centromere_hg38.txt; tr_file=hg38_tandem_repeats.bed;if_hg38=True; pass_only=True; seq_resolved=False; num_X_chr=2; wlen_tp=True
python ttmars.py “$output_dir” “$if_hg38” “$centro_file” assem1_non_cov_regions.bed assem2_non_cov_regions.bed “$vcf_file” “$reference” h1.fa h2.fa lo_pos_assem1_result_compressed.bed lo_pos_assem2_result_compressed.bed “$tr_file” “$pass_only” “$seq_resolved” “$wlen_tp”
python reg_dup.py “$output_dir” “$if_hg38” “$centro_file” assem1_non_cov_regions.bed assem2_non_cov_regions.bed “$vcf_file” “$reference” h1.fa h2.fa lo_pos_assem1_result_compressed.bed lo_pos_assem2_result_compressed.bed “$tr_file” lo_pos_assem1_0_result_compressed.bed lo_pos_assem2_0_result_compressed.bed “$pass_only” “$wlen_tp”
python chrx.py “$output_dir” “$if_hg38” “$centro_file” assem1_non_cov_regions.bed assem2_non_cov_regions.bed “$vcf_file” “$reference” h1.fa h2.fa lo_pos_assem1_result_compressed.bed lo_pos_assem2_result_compressed.bed “$tr_file” “$pass_only” “$seq_resolved” “$wlen_tp”
python combine.py “$output_dir” “$num_X_chr”

For evaluating the genotypes, we calculated non-reference error rates in all trio/duo genotypes by dividing the number of Mendelian-inconsistent genotypes by all sites present in at least one individual of the pedigree. In addition to the error rates, we calculated allelic balances by measuring the ratio between the reference and heterozygous genotypes in children when their parents had reference (REF) and heterozygous (HET) genotypes to identify possible biases in making genotype calls.

### PCA

Ancestries were assigned by identifying the genetic ancestry across individuals^25^. Filtering was performed to exclude variants with AC==1. A randomized PCA was performed using scikit-allel v1.3.5 with n_components=10 and scaler=‘patterson’.

### FST

All variants with AC==1 were excluded. Furthermore, we excluded populations with fewer than 1,000 samples and populations with higher numbers of samples were randomly subsetted to 7,000 samples. Hudson FST was calculated using scikit-allel v1.3.5 (https://zenodo.org/record/4759368#.YbxgLn3MIq0).

### SV-phenotype association in ARIC and HCHS/SOL

The ARIC study is an ongoing biracial cohort designed for cardiovascular research, as described in detail previously^56^. HCHS/SOL is a community-based cohort study of Hispanics/Latinos designed to examine risk and protective factors for chronic diseases, as published previously^57,58^. For our analyses, we selected heart/lung/blood phenotypes related to cardiovascular outcomes that were measured across the entire cohort to maximize sample size. The list of analyzed phenotypes analyzed are height, body mass index (BMI), waist-hip ratio (WHR), whilte blood cell count, hemoglobin, hematocrit, neutrophil, platelet count, systolic/diastolic blood pressure, high-density lipoprotein (HDL), low-density lipoprotein (LDL), triglyceride, high sensitive C-reactive protein (hsCRP), fasting glucose, fasting insulin, glycated hemoglobin (HbA1c), serum creatinine, albumin, albumin-to-creatinine ratio, estimated glomerular filtration rate (eGFR), cystatin C, alanine aminotransferase (ALT), aspartate aminotransferase (AST),and gammaglutamyltransferase (GGT). We tested statistical association of each SV with individual phenotypes separately on ARIC African Americans, European Americans, and HCHS/SOL Hispanics and then performed inverse variance weighted meta-analysis across three populations. We used age, sex, age by sex, age^2^, age^2^ by sex and first five ancestry principal components (PCs) from SNPs and five PCs from SVs as covariates in all analyses. All traits are rank-normalized after adjustment for covariates. We excluded prevalent diabetes cases from glucose and insulin quantitative trait analyses, excluded prevalent CHD cases from hsCRP analyses, and excluded prevalent CKD cases from creatinine, albumin, cystatin C, and albumin to creatinine ratio analyses. Blood pressure levels of hypertension medication users were adjusted by +15 for systolic and +10 for diastolic blood pressure. For ARIC African Americans and European Americans, we used linear regression (Wald) tests for quantitative traits and logistic regression (Wald) tests implemented in the EPACTS pipeline (https://genome.sph.umich.edu/wiki/EPACTS). For HCHS/SOL Hispanic samples, we used GMMAT^59^ package with three random effects for genetic relatedness, household and census block groups to address for population and cohort substructures. Meta analyses were done using METAL software ^60^.

### Linkage disequilibrium (LD) analysis between SNPs and SVs

We first selected SVs and SNPs with minimum minor allele frequency of 0.001 in founders using inferred pedigree information using KING^61^. We also used these founders-only subset in the following analysis. LD calculation was done for SNPs that are within +/- 1Mbp for each SV using PLINK v1.90b6.24 with –r2 –ld-window-r2 0.05 options. We used GWAS catalog downloaded from https://www.ebi.ac.uk/gwas/docs/file-downloads in ‘all associations v1.0’ format on December 13, 2021.

### Annotation of SVs and their impact over genes

We annotated the generated population VCF file using AnnotSV^34^ with default parameters. BCFtools was used to filter and extract SVs that overlapped with genes for different allele frequency (AF=0.05) or allele count (AC=20) thresholds. Subsequently these gene lists were analyzed using DAVID^62^ version 6.8 to identify enrichment of Go terms or KEGG pathways. AnnotSV^63^ v2.5 was run with default parameters, intersecting annotations with gnomad SV (v2.1), and Ensembl Genes (v2021-03-19).

### Hotspots + Valleys

The TOPMed SVs were subset to events ≥50 bp. HGSV SV were downloaded from http://ftp.1000genomes.ebi.ac.uk/vol1/ftp/data_collections/HGSVC2/release/v2.0/integrated_callset/variants_freeze4_sv_insdel_alt.vcf.gz and subsetted to events ≥50 bp. GRCh38 autosomes were split into 100kb disjointed regions with windows intersecting centromere and gap regions as annotated by UCSC genome tracks were removed. The number of SV for each window over the TOPMed variants and HGSV variants were counted. Hotspots were defined as windows with greater than 3x the standard deviation per-variant set. Valleys were defined as windows without any SV per-variant set.

## Data Availability

An overview of the TOPMed participant consents and data access procedures is provided in Taliun et al. ^25^. All TOPMed WGS (cram) data are publicly available on a cloud-based platform with access managed by dbGaP under study specific accessions. The dbGaP accession numbers for all TOPMed studies referenced in this paper are listed in Extended Data Table … and a detailed protocol for data access and a description of the publicly accessible data resources is available at https://topmed.nhlbi.nih.gov/topmed-data-access-scientific-community

Structural variant calls from this joint call set for each cohort will be made available under study specific dbGaP accessions using standardized sample IDs and formats to facilitate combined analyses^25^. The full call set including per sample genotypes is currently available via the TOPMed dbGap Exchange Area for approved TOPMed investigators. To further promote the utilization for this call set we have deposited the SV alleles together with population frequencies in dbVar (accession ID: Jun2023) for studies with appropriate consent. This will allow studies to easily compare their individual SV call sets with ours and utilize the large call set to more robustly annotate their individual SV with population frequencies.

Long-read assemblies for NA12878 and NA19238 used for benchmarking are publicly available at http://ftp.1000genomes.ebi.ac.uk/vol1/ftp/data_collections/HGSVC2/release/v1.0/assemblies/20200628_HHU_assembly-results_CCS_v12/assemblies/phased/v12_NA19238_hgsvc_pbsq2-ccs_1000-pereg.h1-un.racon-p2.fasta http://ftp.1000genomes.ebi.ac.uk/vol1/ftp/data_collections/HGSVC2/release/v1.0/assemblies/20200628_HHU_assembly-results_CCS_v12/assemblies/phased/v12_NA19238_hgsvc_pbsq2-ccs_1000-pereg.h2-un.racon-p2.fasta http://ftp.1000genomes.ebi.ac.uk/vol1/ftp/data_collections/HGSVC2/release/v1.0/assemblies/20200628_HHU_assembly-results_CCS_v12/assemblies/phased/v12_NA12878_giab_pbsq2-ccs_1000-pereg.h1-un.racon-p2.fasta http://ftp.1000genomes.ebi.ac.uk/vol1/ftp/data_collections/HGSVC2/release/v1.0/assemblies/20200628_HHU_assembly-results_CCS_v12/assemblies/phased/v12_NA12878_giab_pbsq2-ccs_1000-pereg.h2-un.racon-p2.fasta

## Code Availability

The variant calling pipelines have been deployed and published previously Parliament2^27^: https://github.com/slzarate/parliament2 MuCNV^29^: https://github.com/gjun/muCNV. Additional analysis scripts are collected here: https://github.com/BCM-HGSC/TopMedSVQC

## Acknowledgements

We would like to acknowledge Joshua Traynelis for his contribution to this study. Furthermore, we would like to thank the entire TOPMed team for constructive discussions and suggestions. A special thanks to Ashley S. Smith for helpful advice and administration.

Genetics of Cardiometabolic Health in the Amish (Amish). The TOPMed component of the Amish Research Program was supported by NIH grants R01 HL121007, U01 HL072515, and R01 AG18728.

The Atherosclerosis Risk in Communities study has been funded in whole or in part with Federal funds from the National Heart, Lung, and Blood Institute, National Institutes of Health, Department of Health and Human Services (contract numbers HHSN268201700001I, HHSN268201700002I, HHSN268201700003I, HHSN268201700004I and HHSN268201700005I). The authors thank the staff and participants of the ARIC study for their important contributions.

Molecular Mechanisms of Inherited Cardiomyopathies and Arrhythmias in the Australian Familial AF Study (AustralianFamilialAF) This study was supported by the Victor Chang Cardiac Research Institute, National Health and Medical Research Council of Australia, Estate of the Late RT Hall, Simon Lee Foundation and the St Vincent’s Clinic Foundation.

New Approaches for Empowering Studies of Asthma in Populations of African Descent - Barbados Asthma Genetics Study (BAGS) We gratefully acknowledge the contributions of Pissamai and Trevor Maul, Paul Levett, Anselm Hennis, P. Michele Lashley, Raana Naidu, Malcolm Howitt and Timothy Roach, and the numerous health care providers, and community clinics and co-investigators who assisted in the phenotyping and collection of DNA samples, and the families and patients for generously donating DNA samples to the Barbados Asthma Genetics Study (BAGS). Funding for BAGS was provided by National Institutes of Health (NIH) R01HL104608, R01HL087699, and HL104608 S1.

The Mount Sinai BioMe Biobank has been supported by The Andrea and Charles Bronfman Philanthropies and in part by Federal funds from the NHLBI and NHGRI (U01HG00638001; U01HG007417; X01HL134588). We thank all participants in the Mount Sinai Biobank. We also thank all our recruiters who have assisted and continue to assist in data collection and management and are grateful for the computational resources and staff expertise provided by Scientific Computing at the Icahn School of Medicine at Mount Sinai.

The Coronary Artery Risk Development in Young Adults Study (CARDIA) is conducted and supported by the National Heart, Lung, and Blood Institute (NHLBI) in collaboration with the University of Alabama at Birmingham (HHSN268201800005I & HHSN268201800007I), Northwestern University (HHSN268201800003I), University of Minnesota (HHSN268201800006I), and Kaiser Foundation Research Institute (HHSN268201800004I). CARDIA was also partially supported by the Intramural Research Program of the National Institute on Aging (NIA) and an intra-agency agreement between NIA and NHLBI (AG0005).

Cleveland Clinic Atrial Fibrillation Study (CCAF) This study was supported by the National Institutes of Health (NIH) grants R01 HL 090620 and R01 HL 111314, the NIH National Center for Research Resources for Case Western Reserve University and Cleveland Clinic Clinical and Translational Science Award UL1-RR024989, the Cleveland Clinic Department of Cardiovascular Medicine philanthropy research funds, and the Tomsich Atrial Fibrillation Research Fund.

Cleveland Family Study - WGS Collaboration (CFS) The Cleveland Family Study has been supported in part by National Institutes of Health grants [R01-HL046380, KL2-RR024990, R35-HL135818, and R01-HL113338].

Children’s Health Study: Integrative Genetic Approaches to Gene-Air Pollution Interactions in Asthma (ChildrensHS_GAP) The Integrative Genetic Approaches to Gene-Air Pollution Interactions in Asthma (GAP) study was supported by the National Institute of Environmental Health Sciences (NIEHS) grant #R01ES021801. The Children’s Health Study (CHS) was supported by the Southern California Environmental Health Sciences Center (grant P30ES007048); National Institute of Environmental Health Sciences (grants 5P01ES011627, ES021801, ES023262, P01ES009581, P01ES011627, P01ES022845, R01 ES016535, R03ES014046, P50 CA180905, R01HL061768, R01HL076647, R01HL087680 and RC2HL101651), the Environmental Protection Agency (grants RD83544101, R826708, RD831861, and R831845), and the Hastings Foundation.

Children’s Health Study: Integrative Genomics and Environmental Research of Asthma (ChildrensHS_IGERA) The Integrative Genomics and Environmental Research of Asthma (IGERA) Study was supported by the National Heart, Lung and Blood Institute (grant # RC2HL101543 -The Asthma BioRepository for Integrative Genomics Research, PI Gilliland/Raby). The Children’s Health Study (CHS) was supported by the Southern California Environmental Health Sciences Center (grant P30ES007048); National Institute of Environmental Health Sciences (grants 5P01ES011627, ES021801, ES023262, P01ES009581, P01ES011627, P01ES022845, R01 ES016535, R03ES014046, P50 CA180905, R01HL061768, R01HL076647, R01HL087680 and RC2HL101651), the Environmental Protection Agency (grants RD83544101, R826708, RD831861, and R831845), and the Hastings Foundation.

Children’s Health Study: Effects of Air Pollution on the Development of Obesity in Children (ChildrensHS_MetaAir) The Effects of Air Pollution on the Development of Obesity in Children (Meta-AIR) study was supported by the Southern California Children’s Environmental Health Center funded by the National Institute of Environmental Health Sciences (NIEHS) (P01ES022845) and the Environmental Protection Agency (EPA) (RD-83544101–0). The Children’s Health Study (CHS) was supported by the Southern California Environmental Health Sciences Center (grant P30ES007048); National Institute of Environmental Health Sciences (grants 5P01ES011627, ES021801, ES023262, P01ES009581, P01ES011627, P01ES022845, R01 ES016535, R03ES014046, P50 CA180905, R01HL061768, R01HL076647, R01HL087680, RC2HL101651 and K99ES027870), the Environmental Protection Agency (grants RD83544101, R826708, RD831861, and R831845), and the Hastings Foundation.

Cardiovascular Health Study: This research was supported by contracts HHSN268201200036C, HHSN268200800007C, HHSN268201800001C, N01HC55222, N01HC85079, N01HC85080, N01HC85081, N01HC85082, N01HC85083, N01HC85086, 75N92021D00006, and grants U01HL080295 and U01HL130114 from the National Heart, Lung, and Blood Institute (NHLBI), with additional contribution from the National Institute of Neurological Disorders and Stroke (NINDS). Additional support was provided by R01AG023629 from the National Institute on Aging (NIA). A full list of principal CHS investigators and institutions can be found at CHS-NHLBI.org. The content is solely the responsibility of the authors and does not necessarily represent the official views of the National Institutes of Health.

The COPDGene project described was supported by Award Number U01 HL089897 and Award Number U01 HL089856 from the National Heart, Lung, and Blood Institute. The content is solely the responsibility of the authors and does not necessarily represent the official views of the National Heart, Lung, and Blood Institute or the National Institutes of Health. The COPDGene project is also supported by the COPD Foundation through contributions made to an Industry Advisory Board comprised of AstraZeneca, Boehringer Ingelheim, GlaxoSmithKline, Novartis, Pfizer, Siemens and Sunovion. A full listing of COPDGene investigators can be found at: http://www.copdgene.org/directory

Diabetes Heart Study (DHS) This work was supported by R01 HL92301, R01 HL67348, R01 NS058700, R01 AR48797, R01 DK071891, R01 AG058921, the General Clinical Research Center of the Wake Forest University School of Medicine (M01 RR07122, F32 HL085989), the American Diabetes Association, and a pilot grant from the Claude Pepper Older Americans Independence Center of Wake Forest University Health Sciences (P60 AG10484).

Evaluation of COPD Longitudinally to Identify Predictive Surrogate End-points (ECLIPSE) The ECLIPSE study (NCT00292552) was sponsored by GlaxoSmithKline. The ECLIPSE investigators included: ECLIPSE Investigators — Bulgaria: Y. Ivanov, Pleven; K. Kostov, Sofia. Canada: J. Bourbeau, Montreal; M. Fitzgerald, Vancouver, BC; P. Hernandez, Halifax, NS; K. Killian, Hamilton, ON; R. Levy, Vancouver, BC; F. Maltais, Montreal; D. O’Donnell, Kingston, ON. Czech Republic: J. Krepelka, Prague. Denmark: J. Vestbo, Hvidovre. The Netherlands: E. Wouters, Horn-Maastricht. New Zealand: D. Quinn, Wellington. Norway: P. Bakke, Bergen. Slovenia: M. Kosnik, Golnik. Spain: A. Agusti, J. Sauleda, P. de Mallorca. Ukraine: Y. Feschenko, V. Gavrisyuk, L. Yashina, Kiev; N. Monogarova, Donetsk. United Kingdom: P. Calverley, Liverpool; D. Lomas, Cambridge; W. MacNee, Edinburgh; D. Singh, Manchester; J. Wedzicha, London. United States: A. Anzueto, San Antonio, TX; S. Braman, Providence, RI; R. Casaburi, Torrance CA; B. Celli, Boston; G. Giessel, Richmond, VA; M. Gotfried, Phoenix, AZ; G. Greenwald, Rancho Mirage, CA; N. Hanania, Houston; D. Mahler, Lebanon, NH; B. Make, Denver; S. Rennard, Omaha, NE; C. Rochester, New Haven, CT; P. Scanlon, Rochester, MN; D. Schuller, Omaha, NE; F. Sciurba, Pittsburgh; A. Sharafkhaneh, Houston; T. Siler, St. Charles, MO; E. Silverman, Boston; A. Wanner, Miami; R. Wise, Baltimore; R. ZuWallack, Hartford, CT. ECLIPSE Steering Committee: H. Coxson (Canada), C. Crim (GlaxoSmithKline, USA), L. Edwards (GlaxoSmithKline, USA), D. Lomas (UK), W. MacNee (UK), E. Silverman (USA), R. Tal-Singer (Co-chair, GlaxoSmithKline, USA), J. Vestbo (Co-chair, Denmark), J. Yates (GlaxoSmithKline, USA). ECLIPSE Scientific Committee: A. Agusti (Spain), P. Calverley (UK), B. Celli (USA), C. Crim (GlaxoSmithKline, USA), B. Miller (GlaxoSmithKline, USA), W. MacNee (Chair, UK), S. Rennard (USA), R. Tal-Singer (GlaxoSmithKline, USA), E. Wouters (The Netherlands), J. Yates (GlaxoSmithKline, USA).

Boston Early-Onset COPD Study (EOCOPD) The Boston Early-Onset COPD Study was supported by R01 HL113264 and U01 HL089856 from the National Heart, Lung, and Blood Institute.

## Framingham Heart Study (FHS)

The Framingham Heart Study (FHS) acknowledges the support of contracts NO1-HC-25195, HHSN268201500001I and 75N92019D00031 from the National Heart, Lung and Blood Institute and grant supplement R01 HL092577-06S1 for this research. We also acknowledge the dedication of the FHS study participants without whom this research would not be possible.

## ATGC Gene-Environment, Admixture and Latino Asthmatics Study I Asthma (GALAI)

The Genes-environments and Admixture in Latino Americans (GALA I) Study was supported by the National Heart, Lung, and Blood Institute of the National Institute of Health (NIH) grants R01HL117004 and X01HL134589; study enrollment supported by Sandler Center for Basic Research in Asthma and the Sandler Family Foundation, the American Asthma Foundation, the American Lung Association, the NIH grants K23HL04464 and HL07185, the Resource Centers for Minority Aging Research from the National Institute on Aging, RCMAR P30-AG15272, the National Institute of Nursing Research and the National Center on Minority Health and Health Disparities.

## Gene-Environment, Admixture and Latino Asthmatics Study (GALAII)

The Genes-environments and Admixture in Latino Americans (GALA II) Study was supported by the National Heart, Lung, and Blood Institute of the National Institute of Health (NIH) grants R01HL117004 and X01HL134589; study enrollment supported by the Sandler Family Foundation, the American Asthma Foundation, the RWJF Amos Medical Faculty Development Program, Harry Wm. and Diana V. Hind Distinguished Professor in Pharmaceutical Sciences II and the National Institute of Environmental Health Sciences grant R01ES015794. The GALA II study collaborators include Shannon Thyne, UCSF; Harold J. Farber, Texas Children’s Hospital; Denise Serebrisky, Jacobi Medical Center; Rajesh Kumar, Lurie Children’s Hospital of Chicago; Emerita Brigino-Buenaventura, Kaiser Permanente; Michael A. LeNoir, Bay Area Pediatrics; Kelley Meade, UCSF Benioff Children’s Hospital, Oakland; William Rodriguez-Cintron, VA Hospital, Puerto Rico; Pedro C. Avila, Northwestern University; Jose R. Rodriguez-Santana, Centro de Neumologia Pediatrica; Luisa N. Borrell, City University of New York; Adam Davis, UCSF Benioff Children’s Hospital, Oakland; Saunak Sen, University of Tennessee and Fred Lurmann, Sonoma Technologies, Inc. The authors acknowledge the families and patients for their participation and thank the numerous health care providers and community clinics for their support and participation in GALA II. In particular, the authors thank study coordinator Sandra Salazar; the recruiters who obtained the data: Duanny Alva, MD, Gaby Ayala-Rodriguez, Lisa Caine, Elizabeth Castellanos, Jaime Colon, Denise DeJesus, Blanca Lopez, Brenda Lopez, MD, Louis Martos, Vivian Medina, Juana Olivo, Mario Peralta, Esther Pomares, MD, Jihan Quraishi, Johanna Rodriguez, Shahdad Saeedi, Dean Soto, Ana Taveras; and the lab researcher Celeste Eng who processed the biospecimens.

## Genetic Epidemiology Network of Arteriopathy (GENOA)

Support for GENOA was provided by the National Heart, Lung and Blood Institute (HL054457, HL054464, HL054481, HL119443, and HL087660) of the National Institutes of Health.

The Genetic Epidemiology Network of Salt-Sensitivity (GenSalt) was supported by research grants (U01HL072507, R01HL087263, and R01HL090682) from the National Heart, Lung and Blood Institute, National Institutes of Health, Bethesda, MD.

The GGAF (Groningen Genetics of Atrial Fibrillation) is supported by funding to the 5 sources that form GGAF. The AF RISK study is supported by the Netherlands Heart Foundation (grant NHS2010B233), and the Center for Translational Molecular Medicine. Both the Young-AF and Biomarker-AF studies are supported by funding from the University Medical Center Groningen. The GIPS-III trial was supported by grant 95103007 from ZonMw, the Netherlands Organization for Health Research and Development. The PREVEND study is supported by the Dutch Kidney Foundation (grant E0.13) and the Netherlands Heart Foundation (grant NHS2010B280).

Genetics of Lipid Lowering Drugs and Diet Network (GOLDN) GOLDN biospecimens, baseline phenotype data, and intervention phenotype data were collected with funding from National Heart, Lung and Blood Institute (NHLBI) grant U01 HL072524. Whole-genome sequencing in GOLDN was funded by NHLBI grant R01 HL104135 and supplement R01 HL104135-04S1.

Hispanic Community Health Study - Study of Latinos (HCHS_SOL) The Hispanic Community Health Study/Study of Latinos is a collaborative study supported by contracts from the National Heart, Lung, and Blood Institute (NHLBI) to the University of North Carolina (HHSN268201300001I / N01-HC-65233), University of Miami (HHSN268201300004I / N01-HC-65234), Albert Einstein College of Medicine (HHSN268201300002I / N01-HC-65235), University of Illinois at Chicago – HHSN268201300003I / N01-HC-65236 Northwestern Univ), and San Diego State University (HHSN268201300005I / N01-HC-65237). The following Institutes/Centers/Offices have contributed to the HCHS/SOL through a transfer of funds to the NHLBI: National Institute on Minority Health and Health Disparities, National Institute on Deafness and Other Communication Disorders, National Institute of Dental and Craniofacial Research, National Institute of Diabetes and Digestive and Kidney Diseases, National Institute of Neurological Disorders and Stroke, NIH Institution-Office of Dietary Supplements.

Heart and Vascular Health Study (HVH) The Heart and Vascular Health Study was supported by grants HL068986, HL085251, HL095080, and HL073410 from the National Heart, Lung, and Blood Institute.

Hypertension Genetic Epidemiology Network (HyperGEN) The HyperGEN Study is part of the National Heart, Lung, and Blood Institute (NHLBI) Family Blood Pressure Program; collection of the data represented here was supported by grants U01 HL054472 (MN Lab), U01 HL054473 (DCC), U01 HL054495 (AL FC), and U01 HL054509 (NC FC). The HyperGEN: Genetics of Left Ventricular Hypertrophy Study was supported by NHLBI grant R01 HL055673 with wholegenome sequencing made possible by supplement −18S1.

The Jackson Heart Study (JHS) is supported and conducted in collaboration with Jackson State University (HHSN268201800013I), Tougaloo College (HHSN268201800014I), the Mississippi State Department of Health (HHSN268201800015I) and the University of Mississippi Medical Center (HHSN268201800010I, HHSN268201800011I and HHSN268201800012I) contracts from the National Heart, Lung, and Blood Institute (NHLBI) and the National Institute on Minority Health and Health Disparities (NIMHD). The authors also wish to thank the staffs and participants of the JHS.

## Mayo Clinic Venous Thromboembolism Study (Mayo_VTE)

Funded, in part, by grants from the National Institutes of Health, National Heart, Lung and Blood Institute (HL66216 and HL83141). the National Human Genome Research Institute (HG04735, HG06379), and research support provided by Mayo Foundation.

Whole genome sequencing (WGS) for the Trans-Omics in Precision Medicine (TOPMed) program was supported by the National Heart, Lung and Blood Institute (NHLBI). WGS for “NHLBI TOPMed: Multi-Ethnic Study of Atherosclerosis (MESA)” (phs001416.v1.p1) was performed at the Broad Institute of MIT and Harvard (3U54HG003067-13S1). Centralized read mapping and genotype calling, along with variant quality metrics and filtering were provided by the TOPMed Informatics Research Center (3R01HL-117626-02S1). Phenotype harmonization, data management, sample-identity QC, and general study coordination, were provided by the TOPMed Data Coordinating Center (3R01HL-120393-02S1), and TOPMed MESA Multi-Omics (HHSN2682015000031/HSN26800004). The MESA projects are conducted and supported by the National Heart, Lung, and Blood Institute (NHLBI) in collaboration with MESA investigators. Support for the Multi-Ethnic Study of Atherosclerosis (MESA) projects are conducted and supported by the National Heart, Lung, and Blood Institute (NHLBI) in collaboration with MESA investigators. Support for MESA is provided by contracts 75N92020D00001, HHSN268201500003I, N01-HC-95159, 75N92020D00005, N01-HC-95160, 75N92020D00002, N01-HC-95161, 75N92020D00003, N01-HC-95162, 75N92020D00006, N01-HC-95163, 75N92020D00004, N01-HC-95164, 75N92020D00007, N01-HC-95165, N01-HC-95166, N01-HC-95167, N01-HC-95168, N01-HC-95169, UL1-TR-000040, UL1-TR-001079, UL1-TR-001420, UL1TR001881, DK063491, and R01HL105756. The authors thank the other investigators, the staff, and the participants of the MESA study for their valuable contributions. A full list of participating MESA investigators and institutes can be found at https://urldefense.proofpoint.com/v2/url?u=http-3A_www.mesa-2Dnhlbi.org&d=DwIGaQ&c=ZQs-KZ8oxEw0p81sqgiaRA&r=hkDaFIlMMg5o5xD5aQ2Txzhw0BLHnUrK9Cxqzcl0pbM&m=o32ACAoh0FIJtL7l5AXiWsB-7bcyn6az6hBBIzImDkB_5H5GFBxgq0ht1DTbWmyc&s=9-Gwf81AIEYr7ywl6zPi7VXkojbPiptVvhZpePkr4Ag&e=. This study was also supported in part by the NHLBI contracts R01HL151855 and R01HL146860.

E.E.E. is an investigator of the Howard Hughes Medical Institute.

My Life, Our Future: Genotyping for Progress in Hemophilia (MLOF) The My Life, Our Future samples and data are made possible through the partnership of Bloodworks Northwest, the American Thrombosis and Hemostasis Network, the National Hemophilia Foundation, and Bioverativ. We gratefully acknowledge the hemophilia treatment centers and their patients who provided biological samples and phenotypic data.

Outcome Modifying Genes in Sickle Cell Disease (OMG_SCD) The OMG-SCD study was administrated by Marilyn J. Telen, M.D. and Allison E. Ashley-Koch, Ph.D. from Duke University Medical Center and collection of the data set was supported by grants HL068959 and HL079915 from the National Heart, Lung, and Blood Institute (NHLBI) of the National Institute of Health (NIH).

## National Institutes of Health (R01HL113326, P30 GM110766-01)

The Pediatric Cardiac Genomics Consortium (PCGC) program is funded by the National Heart, Lung, and Blood Institute, National Institutes of Health, U.S. Department of Health and Human Services through grants UM1HL128711, UM1HL098162, UM1HL098147, UM1HL098123, UM1HL128761, and U01HL131003.

Whole Genome Sequencing to Identify Causal Genetic Variants Influencing CVD Risk - San Antonio Family Studies (SAFS) Collection of the San Antonio Family Study data was supported in part by National Institutes of Health (NIH) grants R01 HL045522, MH078143, MH078111 and MH083824; and whole genome sequencing of SAFS subjects was supported by U01 DK085524 and R01 HL113323. We are very grateful to the participants of the San Antonio Family Study for their continued involvement in our research programs.

Study of African Americans, Asthma, Genes and Environment (SAGE) The Study of African Americans, Asthma, Genes and Environments (SAGE) was supported by by the National Heart, Lung, and Blood Institute of the National Institute of Health (NIH) grants R01HL117004 and X01HL134589; study enrollment supported by the Sandler Family Foundation, the American Asthma Foundation, the RWJF Amos Medical Faculty Development Program, Harry Wm. and Diana V. Hind Distinguished Professor in Pharmaceutical Sciences II. The SAGE study collaborators include Harold J. Farber, Texas Children’s Hospital; Emerita Brigino-Buenaventura, Kaiser Permanente; Michael A. LeNoir, Bay Area Pediatrics; Kelley Meade, UCSF Benioff Children’s Hospital, Oakland; Luisa N. Borrell, City University of New York; Adam Davis, UCSF Benioff Children’s Hospital, Oakland and Fred Lurmann, Sonoma Technologies, Inc. The authors acknowledge the families and patients for their participation and thank the numerous health care providers and community clinics for their support and participation in SAGE. In particular, the authors thank study coordinator Sandra Salazar; the recruiters who obtained the data: Lisa Caine, Elizabeth Castellanos, Brenda Lopez, MD, Shahdad Saeedi; and the lab researcher Celeste Eng who processed the biospecimens.

The Samoan Obesity, Lifestyle and Genetic Adaptations Study (OLaGA) Group’. In any supplementary files please list the names in the OLAGA group, as follows: Ranjan Deka, Dept. of Environmental Health, University of Cincinnati; Nicola L. Hawley, Dept. of Chronic Disease Epidemiology, Yale University; Stephen T McGarvey, Dept. of Epidemiology and International Health Institute, and Dept. of Anthropology, Brown University; Ryan L Minster, Dept. of Human Genetics, University of Pittsburgh; Take Naseri, Ministry of Health, Government of Samoa; Muagututi‘a Sefuiva Reupena, Lutia I Puava Ae Mapu I Fagalele; Daniel E. Weeks, Depts. of Human Genetics and Biostatistics, University of Pittsburgh.

Study of African Americans, Asthma, Genes and Environment (SAGE) The Study of African Americans, Asthma, Genes and Environments (SAGE) was supported by by the National Heart, Lung, and Blood Institute of the National Institute of Health (NIH) grants R01HL117004 and X01HL134589; study enrollment supported by the Sandler Family Foundation, the American Asthma Foundation, the RWJF Amos Medical Faculty Development Program, Harry Wm. and Diana V. Hind Distinguished Professor in Pharmaceutical Sciences II. The SAGE study collaborators include Harold J. Farber, Texas Children’s Hospital; Emerita Brigino-Buenaventura, Kaiser Permanente; Michael A. LeNoir, Bay Area Pediatrics; Kelley Meade, UCSF Benioff Children’s Hospital, Oakland; Luisa N. Borrell, City University of New York; Adam Davis, UCSF Benioff Children’s Hospital, Oakland and Fred Lurmann, Sonoma Technologies, Inc. The authors acknowledge the families and patients for their participation and thank the numerous health care providers and community clinics for their support and participation in SAGE. In particular, the authors thank study coordinator Sandra Salazar; the recruiters who obtained the data: Lisa Caine, Elizabeth Castellanos, Brenda Lopez, MD, Shahdad Saeedi; and the lab researcher Celeste Eng who processed the biospecimens.

Genetics of Sarcoidosis in African Americans (Sarcoidosis) National Institutes of Health (R01HL113326, P30 GM110766-01)

The Rare Variants for Hypertension in Taiwan Chinese (THRV) is supported by the National Heart, Lung, and Blood Institute (NHLBI) grant (R01HL111249) and its participation in TOPMed is supported by an NHLBI supplement (R01HL111249-04S1). THRV is a collaborative study between Washington University in St. Louis, LA BioMed at Harbor UCLA, University of Texas in Houston, Taichung Veterans General Hospital, Taipei Veterans General Hospital, Tri-Service General Hospital, National Health Research Institutes, National Taiwan University, and Baylor University. THRV is based (substantially) on the parent SAPPHIRe study, along with additional population-based and hospital-based cohorts. SAPPHIRe was supported by NHLBI grants (U01HL54527, U01HL54498) and Taiwan funds, and the other cohorts were supported by Taiwan funds.

Treatment of Pulmonary Hypertension and Sickle Cell Disease with Sildenafil Therapy (walk_PHaSST) We thank Dr. Mark Gladwin and the investigators of the Walk-PHasst study and the patients who participated in the study. We also thanks the walk-PHaSST clinical site team: Albert Einstein College of Medicine: Jane Little and Verlene Davis; Columbia University: Robyn Barst, Erika Rosenzweig, Margaret Lee and Daniela Brady; UCSF Benioff Children’s Hospital Oakland: Claudia Morris, Ward Hagar, Lisa Lavrisha, Howard Rosenfeld, and Elliott Vichinsky; Children’s Hospital of Pittsburgh of UPMC: Regina McCollum; Hammersmith Hospital, London: Sally Davies, Gaia Mahalingam, Sharon Meehan, Ofelia Lebanto, and Ines Cabrita; Howard University: Victor Gordeuk, Oswaldo Castro, Onyinye Onyekwere, Vandana Sachdev, Alvin Thomas, Gladys Onojobi, Sharmin Diaz, Margaret Fadojutimi-Akinsiku, and Randa Aladdin; Johns Hopkins University: Reda Girgis, Sophie Lanzkron and Durrant Barasa; NHLBI: Mark Gladwin, Greg Kato, James Taylor, Wynona Coles, Catherine Seamon, Mary Hall, Amy Chi, Cynthia Brenneman, Wen Li, and Erin Smith; University of Colorado: Kathryn Hassell, David Badesch, Deb McCollister and Julie McAfee; University of Illinois at Chicago: Dean Schraufnagel, Robert Molokie, George Kondos, Patricia Cole-Saffold, and Lani Krauz; National Heart & Lung Institute, Imperial College London: Simon Gibbs. Thanks also to the data coordination center team from Rho, Inc.: Nancy Yovetich, Rob Woolson, Jamie Spencer, Christopher Woods, Karen Kesler, Vickie Coble, and Ronald W. Helms. We also thank Dr. Yingze Zhang for directing the Walk-PHasst repository and Dr. Mehdi Nouraie for maintaining the Walk-PHasst database and Dr. Jonathan Goldsmith as a NIH program director for this study. Special thanks to the volunteers who participated in the Walk-PHaSST study. This project was funded with federal funds from the NHLBI, NIH, Department of Health and Human Services, under contract HHSN268200617182C. This study is registered at www.clinicaltrials.gov as NCT00492531. Detail description of the study was published in Blood, 2011 118:855-864, Machado et al “Hospitalization for pain in patients with sickle cell disease treated with sildenafil for elevated TRV and low exercise capacity”.

Women’s Genome Health Study (WGHS) The WGHS is supported by the National Heart, Lung, and Blood Institute (HL043851 and HL080467) and the National Cancer Institute (CA047988 and UM1CA182913). The most recent cardiovascular endpoints were supported by ARRA funding HL099355.

The WHI program is funded by the National Heart, Lung, and Blood Institute, National Institutes of Health, U.S. Department of Health and Human Services through contracts 75N92021D00001, 75N92021D00002, 75N92021D00003, 75N92021D00004, 75N92021D00005.

Molecular data for the Trans-Omics in Precision Medicine (TOPMed) program was supported by the National Heart, Lung and Blood Institute (NHLBI). See the TOPMed Omics Support Table for study specific omics support information. Core support including centralized genomic read mapping and genotype calling, along with variant quality metrics and filtering were provided by the TOPMed Informatics Research Center (3R01HL-117626-02S1; contract HHSN268201800002I). Core support including phenotype harmonization, data management, sample-identity QC, and general program coordination were provided by the TOPMed Data Coordinating Center (R01HL-120393; U01HL-120393; contract HHSN268201800001I). We gratefully acknowledge the studies and participants who provided biological samples and data for TOPMed.

